# A Xeno-free Media for the In Vitro Expansion of Human Spermatogonial Stem Cells

**DOI:** 10.1101/2021.06.04.447118

**Authors:** Meghan Robinson, Luke Witherspoon, Stephanie Willerth, Ryan Flannigan

**Affiliations:** Vancouver Prostate Centre, 2660 Oak St, Vancouver, British Columbia, Canada V6H 3Z6; Department of Urologic Sciences, University of British Columbia, 2329 West Mall, Vancouver, British Columbia, Canada V6T 1Z4; Department of Urology, The Ottawa Hospital, 501 Smyth Rd, Ottawa, Ontario, Canada K1H 8L6; Division of Medical Sciences, University of Victoria, 3800 Finnerty Road, Victoria, British Columbia, Canada V8P 5C2; Department of Mechanical Engineering, University of Victoria, Victoria, British Columbia, Canada; School of Biomedical Engineering, University of British Columbia, 2329 West Mall, Vancouver, British Columbia, Canada V6T 1Z4; Department of Urology, Weill Cornell Medicine, 1300 York Ave, New York, New York 10065, United States

**Keywords:** Adult Germline Stem Cells, Cell Culture Techniques, Humans, Insulin-Like Growth Factor 1, Prostaglandins

## Abstract

*In vitro* expansion of spermatogonial stem cells (SSCs) has been established using animal-derived fetal bovine serum (FBS) and bovine serum albumin (BSA). However, the use of animal components during cell culture introduces the risk of contaminating cells with pathogens, and leads to animal epitope expression, rendering them unsuitable for medical use. Therefore, this study set out to develop a xeno-free, fully defined media for the expansion of human SSCs. We show that the molecules Prostaglandin D2 (PGD-2) and Insulin-Like Growth Factor 1 (IGF-1) can replace FBS and BSA in cell culture media without loss of viability or expansion capability, and that Rho-Associated, Coiled-Coil Containing Protein Kinase (ROCK) inhibitor Y-27632 supplementation improves viability after cryopreservation. Long-term SSC cultures expanded in xeno-free, defined culture conditions shared identical protein expression profiles for well-known SSC markers, while gene expression analyses revealed a significant improvement in quiescent SSC and pan-germ markers. This xeno-free, defined formulation allows for standardized SSC culture free of animal pathogens.

## 1.0 INTRODUCTION^1^

Oncological therapies involving radiation and chemotherapy can be severely detrimental to male fertility, with fertility preservation becoming increasingly highlighted as a key component of cancer survivorship programs.[1] However, many patients undergoing cancer therapies are unable to access the traditional methods of fertility preservation. This is certainly true in the pre-pubertal pediatric population where sperm banking is not possible and cryopreservation of testis tissue remains in the experimental phases of use.[2, 3] Although xenotransplantation of spermatogonial stem cells (SSCs) and testis tissue has been explored, success has been limited, and concerns about possible retrovirus transmission remain.[4–7] *In vitro* systems for restoring spermatogenesis have proven successful using neonatal mice testicular tissues,[8, 9] but have yet to be optimized for human testicular tissues due to a lack of available tissue for research.[10] Orthotopic auto-transplantation of SSCs and testicular tissue in animal models have shown promise, with complete recovery of spermatogenesis, but methods to exclude the risk of re-introducing cancer cells are needed.[10–12] SSCs may represent the most promising therapeutic avenue, ultimately allowing autologous transplantation, or even *in vitro* spermatogenesis.[10, 13] SSCs have been successfully isolated from prepubertal testicular tissues,[14] and shown to expand and maintain a stable epigenetic profile *in vitro*. [15] Furthermore, SSC autologous transplantation studies using irradiated macaques rendered infertile by alkylating or irradiating chemotherapy showed that they are capable of restoring spermatogenesis.[10, 16, 17] While re-introducing cancer cells remains a risk with SSC autologous transplantation, culture methods and sorting strategies are being actively investigated to remove contaminating cancer cells from SSC cultures,[13] and in one promising pilot study, co-culture of acute lymphoblastic leukemia cells with SSCs for two weeks was found to eliminate cancer cells from three different cancer patients as evidenced by highly sensitive polymerase chain reaction to identify the presence of leukemic cells.[18]

A critical step towards adoption of SSC transplantation to restore fertility is refinement of the *in vitro* expansion process.[10, 19] It is estimated that SSC populations will require a several hundred fold increase to achieve adequate transplantation cell volumes,[14] making appropriate cell culture techniques an avenue that must be explored to ensure safe culture conditions. Current cell culture techniques often rely heavily on animal derived additives to help drive cell population expansion and growth.[20] However, this long-standing approach is coming under increased scrutiny given that animal derived growth factors may alter typical cell gene and protein expression from that found in normal physiological *in vivo* conditions,[21] even causing expression of animal epitopes,[22] while variation between batches of animal derived growth factors limits the standardization of cell culture protocols.[23, 24] Furthermore, concerns exist as to the possibility of animal derived antigens and infectious agents that could be transmitted into a human, should *in vitro* expanded cells be transplanted back into human beings.[25–28] Limited research has gone into the production of animal serum free media to help address this issue, with most work focused on mesenchymal derived stem cells and pluripotent stem cells.[29–31] Currently, standard SSC expansion media is composed of Stempro34^™^ SFM, a Food and Drug Administration (FDA)-approved xeno-free, defined basal media[32], supplemented with additional additives including the undefined animal-derived components fetal bovine serum (FBS) and bovine serum albumin (BSA).[14, 33, 34] We show that the addition of the spermatogenic growth factors Insulin-Like Growth Factor-1 (IGF-1), an SSC proliferation regulator,[35] and Prostaglandin D2 (PGD-2), an SSC quiescence regulator,[36–38] can replace the animal serum components in this formulation to enable xeno-free *in vitro* expansion of SSCs, as a step towards translation to clinical use. This study provides a novel protocol for xeno-free expansion of SSCs, and presents comparative phenotype analyses using reverse transcription quantitative polymerase chain reaction (RT-qPCR) and immunocytochemistry analyses of primary SSCs expanded using a xeno-free, chemically defined media versus standard animal serum-supplemented media.

## 2.0 METHODS

### 2.1 Ethical Approval

Testis biopsy samples were obtained through the University of British Columbia Andrology Biobank, with informed consent for research (CREB approved protocol H18-03543). Experiments using hiPSCs in this study were not subject to ethics approval from the University of British Columbia Clinical Research Ethics Boards or Stem Cell Oversight Committee, since they were derived from somatic cells and not intended for transfer into humans or non-human animals.

### 2.2 Human induced pluripotent stem cell (hiPSC) culture

The hiPSC line used was the 1-DL-01 line from WiCell.[39] hiPSCs were expanded on Growth Factor Reduced Matrigel (Corning, 354230) in mTeSR^™^-Plus medium (STEMCELL Technologies, 100-0276), at 37°C and 5%CO_2_, and passaged when 90% confluent using ReLeSR^™^ enzyme-free selective passaging reagent (STEMCELL Technologies, 05872) to maintain purity.

### 2.3 Differentiation of SSCs from hiPSCs (hSSCs)

hSSCs were differentiated from hiPSCs as previously described,[40] with a magnetic activated cell sorting (MACS) step to purify for Thy-1 Cell Surface Antigen (THY1/CD90)-expressing cells. hiPSCs were grown on Matrigel in Minimum Essential Medium Alpha (αMEM, Gibco A10490-01), 1X GlutaMAX (Thermofisher, 35050079), 1X Insulin Transferrin Selenium Liquid Media Supplement (ITS, Millipore Sigma, I13146), 0.2% BSA (Miltenyi Biotec, 130-091-376), 1% penicillin/streptomycin (Sigma Aldrich, P4333), 1 ng/mL human recombinant Fibroblast Growth Factor 2 (FGF2, STEMCELL Technologies, 78003.1), 20 ng/mL animal component-free human recombinant Glial Cell Derived Neurotrophic Factor (GDNF, STEMCELL Technologies, 78139), 0.2% Chemically Defined Lipid Concentrate (Thermofisher, 11905031), and 200 μg/mL L-ascorbic acid (Sigma, A4544). Medium was changed every 2 days for 12-15 days and cells were grown at 37°C and 5% CO_2_. Cells were then switched to expansion medium as previously described,[33] composed of StemPro-34^™^ SFM (Thermofisher, 10639011) supplemented with 1X ITS, 30 μg/ml sodium pyruvate (Sigma, S8636), 1 μl/mL sodium DL-lactic acid solution (Sigma, L4263), 5 mg/ml BSA, 1% FBS (Gibco, 12483-020), 1X GlutaMAX, 5 × 10^-5^ M 2-mercaptoethanol (Gibco, 31350-010), 1X Minimal Essential Medium (MEM) Vitamin Solution (Gibco, 11120052), 10^-4^ M L-ascorbic acid, 10 μg/ml biotin (Sigma, B4639), 30 ng/ml β-estradiol (Sigma, E2758), 60 ng/ml progesterone (Sigma, P8783), 20 ng/mL human recombinant Epidermal Growth Factor (EGF, STEMCELL Technologies, 78006.1), 10 ng/mL human recombinant Leukemia Inhibitory Factor (LIF, STEMCELL Technologies, 78055.1), and 10 ng/mL GDNF. hSSCs were passaged onto CellAdhere^™^ Laminin-521 coated plates (STEMCELL Technologies, 77004) using TrypLE^™^ Express Enzyme (Thermofisher, 12604013). 12 million cells were purified for CD90^+^ cells using a PE-conjugated antibody for anti-CD90 (R&D Systems, FAB2067P), the EasySep^™^ Human PE Positive Selection Kit II (STEMCELL Technologies, 17664), and EasySep^™^ Magnet (STEMCELL Technologies, 18000).

### 2.4 Primary SSC isolation and culture

Primary testicular cells were isolated from human testicular biopsies as previously described.[41] Testicular biopsies were transported to the lab on ice in Hypothermosol^®^ FRS (STEMCELL Technologies, 07935) and processed within 1-2 hours. They were rinsed three times with Hank’s Balanced Salt Solution (HBSS, Millipore Sigma, 55021C), cut into 1mm^3^ pieces with surgical scissors, and digested by Collagenase NB4 (Nordmark Biochemicals, S1745402) at 2 PZU / 100 mg tissue, for 5 minutes at 37°C and 250 rpm. The sample was vigorously shaken and incubated for a further 3 minutes at 37°C and 250 rpm. After spinning for 5 minutes at 200 x g, sedimented tubules were rinsed 3 times with HBSS, with spinning at 200 x g for 5 minutes in between each rinse. The sedimented tubules were re-suspended in 0.25% Trypsin/EDTA (Sigma, T3924) with 0.8 kU / 100mg DNase I (Sigma, D4263), and pipetted 3-5 times with a 5 mL pipette followed by incubation for 5 minutes at 37°C. This process of pipetting followed by incubation was repeated for a total of 3 incubations. FBS was added to stop the digestion at a final concentration of 10%, and the tubules were filtered through a 70 μm filter, followed by a 40μm filter, and centrifuged for 15 minutes at 600 x g. The cells were then sorted into somatic and germ fractions by overnight plating in a single well of a tissue culture-treated 6-well plate in media as described for hSSC culture. Non-adherent cells (germ cells) were transferred the next day to a single well of a CellAdhere^™^ laminin 521-coated 6-well plate, and expanded in the same media at 37°C and 5% CO_2_.

### 2.5 Experimental conditions for xeno-free expansion

hSSCs were used in place of primary SSCs for preliminary experiments to minimize the loss of human primary SSCs, which are not often available for research. Human recombinant IGF-1 (Peprotech, 100-11) and the small molecule PGD-2 (Peprotech, 4150768) were added to cultures of hiPSC-derived SSCs in SSC expansion media without BSA and FBS, separately or in combination, at 10 ng/mL each. The concentration was chosen to match that of the other growth factors LIF, GDNF and FGF-2 already present in the expansion media. Cell viability and expandability was assessed over the course of a single passage grown at 37°C and 5% CO_2_ and compared to a control group in the standard media. Viability was also assessed after cryopreservation and thawing, with or without 10 ng/mL ROCK inhibitor Y-27632 (STEMCELL Technologies, 72302). The condition which promoted the best viability and propagation was then applied to primary SSC cultures for 3 passages at 37°C and 5% CO_2_ and 2 freeze-thaw cycles. Cells were cryopreserved in serum-free, chemically defined CryoStor^™^ CS10 (STEMCELL Technologies, 07930), placed in a Mr. Frosty^™^ (Nalgene) cooling container at −80°C for 24 hours, followed by transfer to liquid nitrogen. Cells were imaged using an EVOS XL Core imaging System (Invitrogen), and images were processed with ImageJ open source software.

### 2.6 Immunocytochemistry

Cells were fixed for 15 minutes in 4% paraformaldehyde solution (PFA, Thermo Scientific, J19943-K2), permeabilized for 15 minutes in 0.1% Triton X-100 (Sigma, X100) in phosphate buffered saline (PBS), and blocked for 2 hours in 5% normal goat serum (NGS, Abcam, ab7481) in PBS. Primary antibodies were diluted in PBS as follows: anti-Octamer-Binding Protein 4 (OCT4, Abcam, ab184665) 1:500, anti-SRY-Box Transcription Factor 9 (SOX9, Abcam, ab76997) 1:500, anti-Thy-1 Cell Surface Antigen (THY1/CD90, Abcam, ab133350) 1:200, anti-GDNF Family Receptor Alpha 1 (GFRA1, Abcam, ab84106) 1:200, anti-Stage-Specific Embryonic Antigen-4 (SSEA4, Abcam ab16287) 1:300, anti-G Protein-Coupled Receptor 125 (GPR125, Abcam, ab51705) 1:200, anti-Actin Alpha 2, Smooth Muscle (ACTA2, Thermofisher, 14-9760-82) 1:500, and incubated overnight at 4°C in the dark. Cells were rinsed 3 times with PBS for 15 minutes each at 4°C in the dark. Goat anti-Rabbit IgG (H+L) Highly Cross-Adsorbed Secondary Antibody Alexa Fluor 488 (Thermofisher, A-11034) or Goat anti-Mouse IgG (H+L) Highly Cross-Adsorbed Secondary Antibody Alexa Fluor 568 (Thermofisher, A-11031) were diluted 1:200 in PBS and incubated with the cells for 4 hours at 4°C in the dark. Cells were rinsed another 3 times with PBS for 15 minutes each at 4°C in the dark. 4’,6-diamidino-2-phenylindole (DAPI, Abcam, ab228549) was diluted to 2.5 μM in PBS and added to the cells for 15 minutes in the dark at room temperature, and replaced by PBS. Cells were imaged using a Zeiss AXio Observer microscope equipped with laser excitation and fluorescence filters for AlexaFluor 488, AlexaFluor 568 and DAPI, and images were processed using ZEN Blue and ImageJ open source software.

### 2.7 Reverse Transcription Qualitative Polymerase Chain Reaction (RT-qPCR)

RNA was extracted using an RNeasy Plus Micro Kit (Qiagen, 74034), and checked for integrity using an Agilent 2200 Tapestation System with High Sensitivity RNA Screentape (Agilent, 5067-5579), High Sensitivity RNA ScreenTape Sample Buffer (Agilent, 5067-5580), and High Sensitivity RNA ScreenTape Ladder (Agilent, 5067-5581). cDNA was generated using iScript^™^ Reverse Transcription Supermix (Bio-Rad, 1708840) with a Tetrad2 Peltier Thermal Cycler (Bio-Rad). RT-qPCR was done with SsoAdvanced^™^ Universal SYBR^®^ Green Supermix (BioRad, 1725270) on a LightCycler96 (Roche). Primers used were PrimePCR^™^ SYBR^®^ Green Assays as listed in **Table 1**. Technical replicates were carried out in triplicate. Analyses was done in Excel and GraphPad Prism Software. Ct values were normalized to Glyceraldehyde-3-Phosphate Dehydrogenase (GAPDH). Technical replicate outliers were detected using Grubbs’ Test, with α=0.05. The 3^rd^ control biological replicate was excluded as an outlier. Results of RT-qPCR are presented as the average Relative Quantification (RQ=2^-ΔΔCt^) values and standard deviations of the biological replicates. Any undetected samples were given a Ct value of the maximum detected cycles plus 1.

**Table 1.**
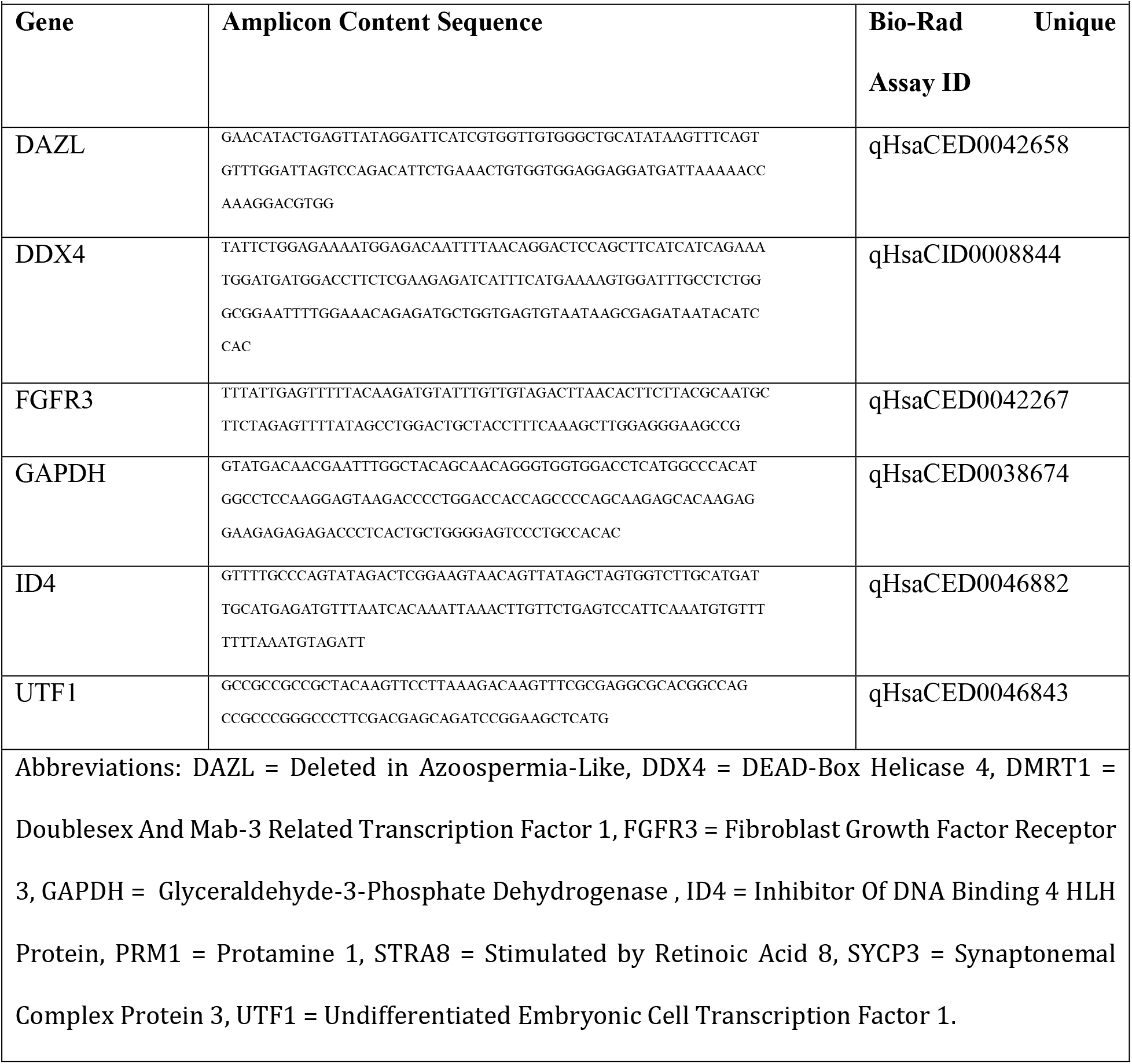
Amplicon content sequences and Unique Assay IDs for Bio-Rad PrimePCR^™^ primer pairs used in real time quantitative polymerase chain reaction (RT-qPCR) assays. Amplicon content sequences (amplicon sequence with additional base pairs added to the beginning and/or end of the sequence), and Unique Assay IDs for Bio-Rad PrimePCR^™^ primer pairs used for RT-qPCR.

### 2.8 Statistics

Statistics were performed using GraphPad Prism software. Each experiment was performed in biological triplicate. Significance between groups was determined by comparing ΔCt values using a student’s unpaired two-tailed t-test, with α=0.05.

See KRT for further information on the essential materials and resources described above.

## 3.0 RESULTS

Because of the rare availability of human testicular tissue for research, we derived SSCs from hiPSCs (hSSCs) for preliminary experimentation of the xeno-free media formulation, before experimenting on primary SSCs over long-term cultures.

### 3.1 Characterization of hSSCs and primary SSCs

After expanding isolated primary SSCs (**Figure 1A**) and differentiated hSSCs (**Figure 1B**), we validated their phenotypes by immunocytochemistry. Undifferentiated hiPSCs showed positive immunoreactivity for the pluripotency marker OCT4 as expected, while hSSCs were negative, confirming a loss of pluripotency (**Figure 1C**). hSSCs were also positive for the well-known SSC markers GPR125, GFRA1, CD90/THY1, and SSEA4, confirming their successful adoption of an SSC-like phenotype (**Figure 1C**). Likewise, primary SSCs showed positive immunoreactivity for GPR125, GFRA1, CD90 and SSEA4, and were negative for the testicular somatic cell markers ACTA2 and SOX9, confirming there was no contamination with somatic cell types (**Figure 1D**).

**Figure 1:**
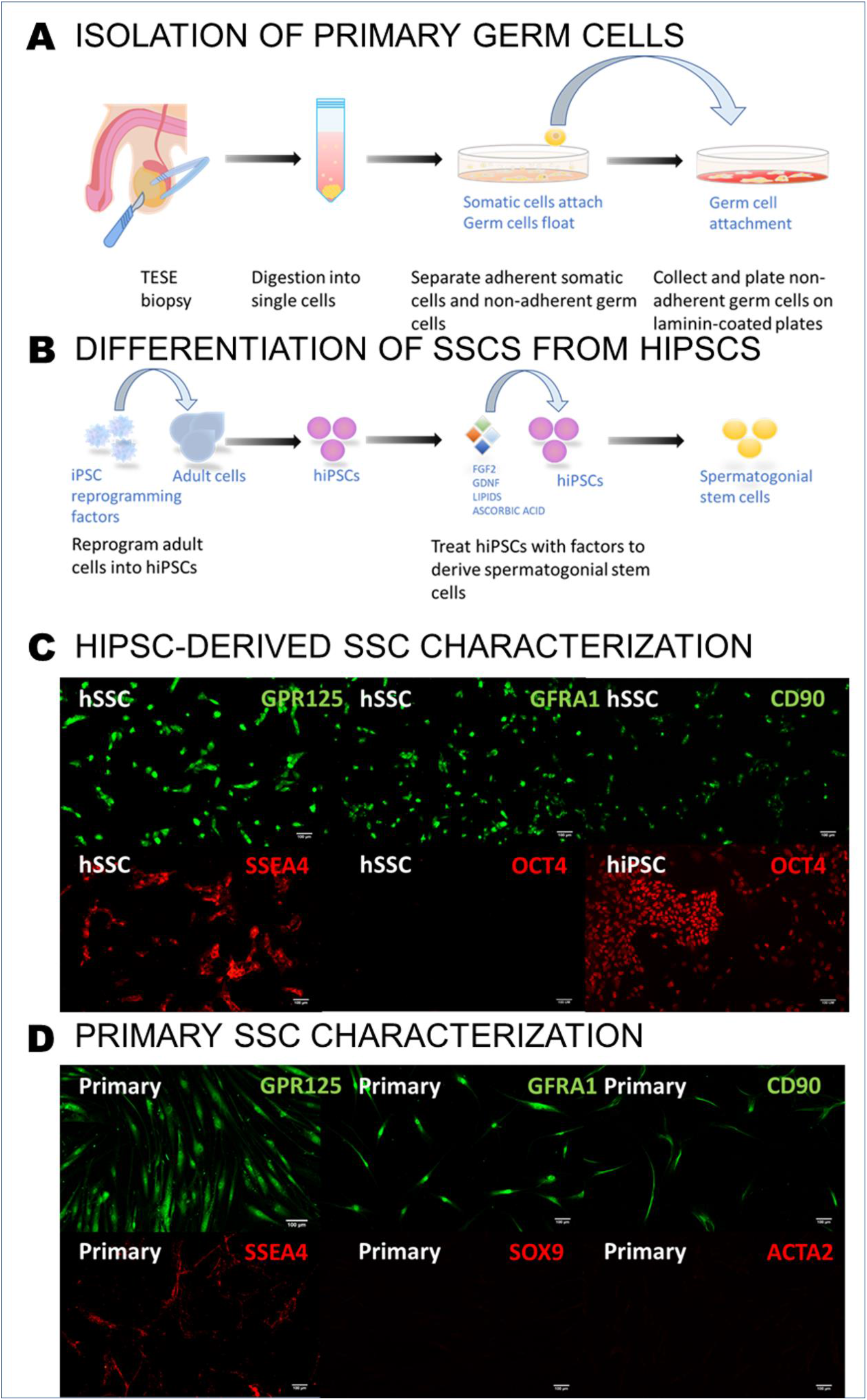
Isolation/derivation and characterization of primary SSCs and hSSCs. **A)** Schematic representing primary SSC isolation: Testicular Sperm Extraction (TESE) biopsies were dissociated into single cells, plated on tissue culture plates, and then 24 hours later the non-adherent germ cells were collected and plated on laminin-coated plates. **B)** Schematic representing SSC derivation from hiPSCs: hiPSCs are reprogrammed from adult fibroblasts and then cultured with FGF2, GDNF, lipids, and ascorbic acid to promote SSC differentiation. **C)** hSSC expression of SSC markers GPR125, GFRA1, CD90 and SSEA4, and the pluripotency factor OCT4, and hiPSC expression of OCT4. **D)** Primary SSC expression of the SSC markers GPR125, GFRA1, CD90 and SSEA4, and the testicular somatic markers ACTA2 and SOX9. All scale bars are 100 μm. **Abbrevations:** FGF2 = Fibroblast Growth Factor 2, GDNF = Glial Cell-Derived Neurotrophic Factor, GPR125 = G-Protein Coupled Receptor 125, GFRA1 = GDNF Family Receptor Alpha 1, CD90 = Thy-1 Cell Surface Antigen, SSEA4 = Stage-Specific Embryonic Antigen 4, OCT4 = Octamer Binding Protein 4, ACTA2 = Actin Alpha 2, Smooth Muscle, SOX9 = SRY=Box Transcription Factor 9, hiPSC = human induced pluripotent stem cell, SSC = spermatogonial stem cell, hSSC = human induced pluripotent stem cell-derived spermatogonial stem cell, Primary = primary spermatogonial stem cell, TESE = testicular sperm extraction.

### 3.2 Establishment of xeno-free media

Removal of the animal components FBS and BSA from standard SSC expansion media induced cell death, therefore we experimented with the supplementation of two spermatogenic factors: IGF-1 and PGD-2 (**Figure 2A**).[35, 36] We found that the addition of both was sufficient to rescue the cells (**Figure 2C**). Furthermore, these factors promoted viability and growth of the hSSCs over several passages on par with the standard media condition (**Figure 2B)**. Interestingly, IGF-1 or PGD-2 alone could not rescue the cells (**Figure 2C**).

**Figure 2:**
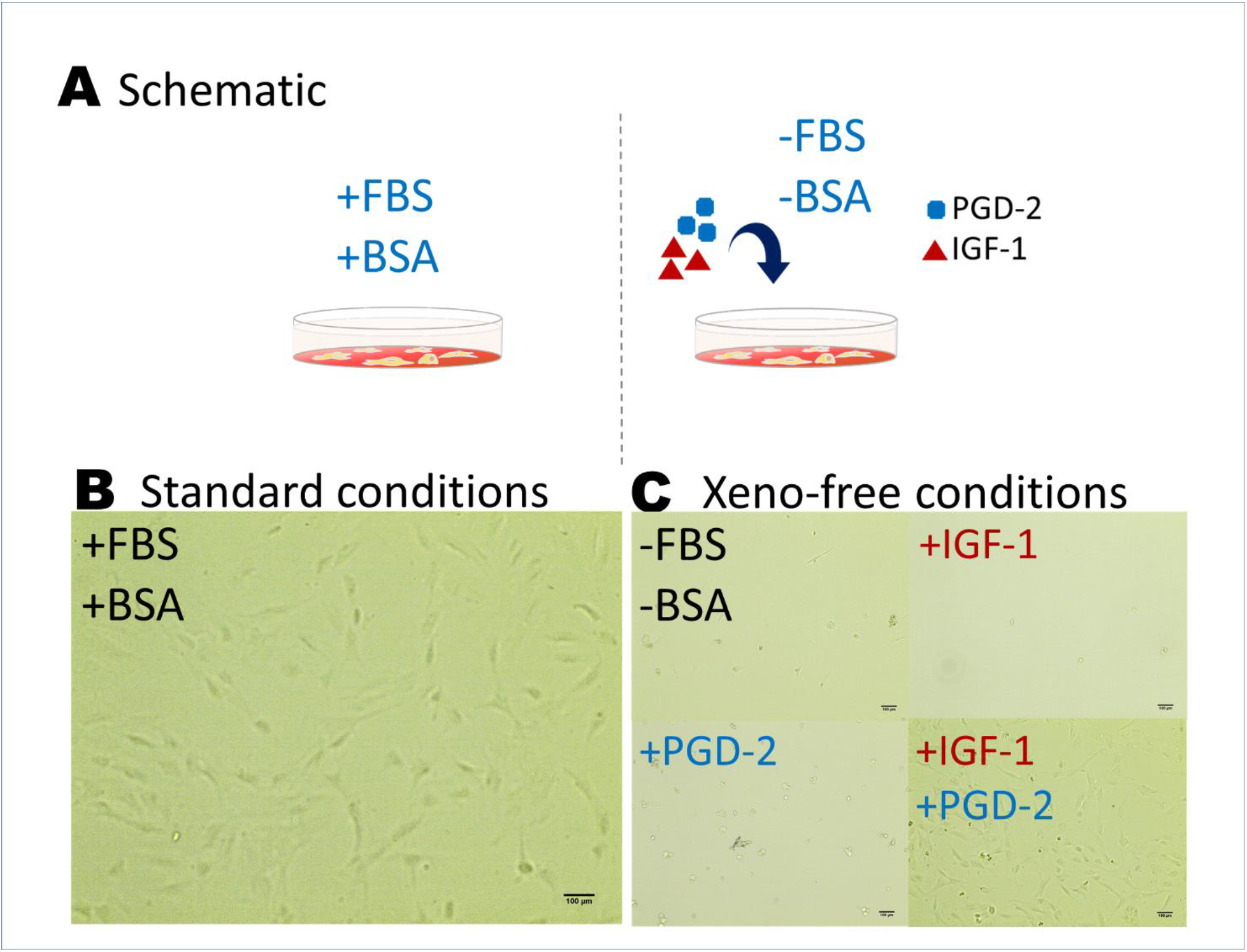
Survival of hSSCs in xeno-free media conditions. **A)** Schematic representing the experimental culture conditions: the standard conditions included FBS and BSA supplementation, while the experimental xeno-free conditions had removed FBS and BSA and additional supplementation of PGD-2 and IGF-1. **B)** Microscope image of the hSSCs in standard conditions. **C)** Microscope images of the hSSCs in the experimental xeno-free media without IGF-1 or PGD-2, with IGF-1 or PGD-2, and with both IGF-1 and PGD-2. All scale bars are 100 μm. **Abbreviations:** hSSC = human induced pluripotent stem cell-derived spermatogonial stem cell, FBS = fetal bovine serum, BSA = bovine serum albumin, PGD-2 = prostaglandin D2, IGF-1 = Insulin-Like Factor 1.

Cryopreserved hSSCs were not viable upon thawing in the xeno-free media formulation, but could be rescued by the one-time addition of the apoptosis inhibitor Y-27632[42] during seeding (**Figure 3B**), and remained viable and expandable thereafter.

**Figure 3:**
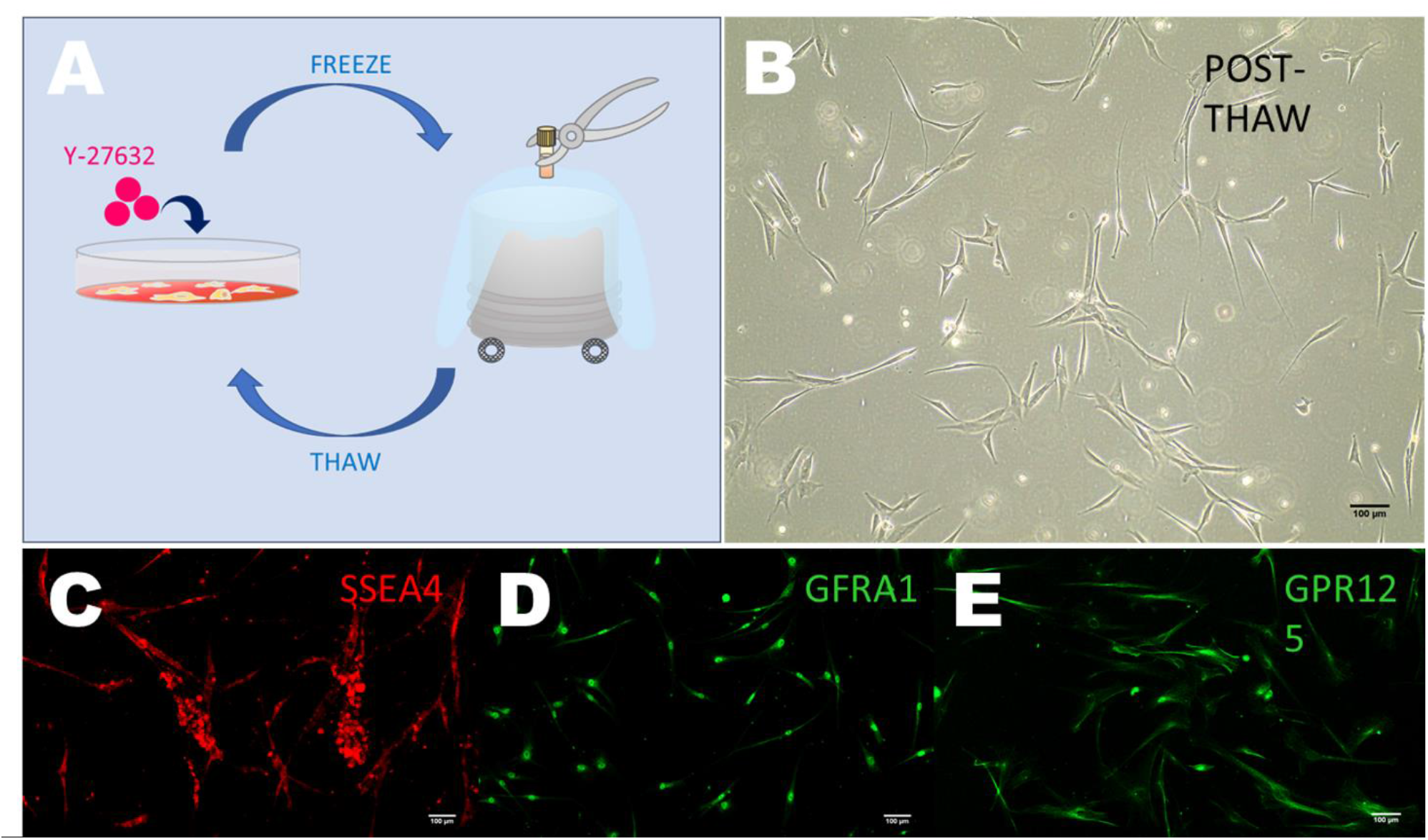
Long-term culture and characterization of primary SSCs in xeno-free conditions. **A)** Schematic representing long-term culture and freeze-thaw cycles before characterization: primary SSCs were expanded, frozen in liquid nitrogen, and then thawed with Y-27632 to promote viability, for a total of 3 cycles, or a 216-fold expansion. **B)** Microscope image of the primary SSCs 24 hours post-thaw in xeno-free conditions. **C-E)** Expression of the SSC markers SSEA4, GFRA1 and GPR125 after 216-fold expansion and 3 freeze-thaw cycles in xeno-free conditions. All scale bars are 100 μm. **Abbreviations:** SSC = spermatogonial stem cell, SSEA4 = Stage-Specific Embryonic Antigen-4, GFRA1 = GDNF Family Receptor Alpha 1, GPR125 = G Protein-Coupled Receptor 125.

### 3.3 Comparison of primary SSCs phenotypes in xeno-free media compared to standard media

Human primary SSCs were grown to confluence in 6-well plates, cryopreserved, and then thawed at a ratio of one cryopreserved well into a fresh 6-well plate (1:6). After three freeze-thaw cycles (roughly 3-4 weeks in culture and a 216-fold expansion, **Figure 3A**), primary SSCs grown in the xeno-free media formulation retained expression of the SSC markers GFRA1, CD90, and SSEA4 (**Figure 3C-E**).[43, 44] Over time, they also began to form grape-like clusters (**Figure 3C**), which is a morphology more typical of SSCs cultured on feeder cells.[45–48]

Gene analyses revealed no significant differences in gene expression between xeno-free and standard media for the quiescent SSC markers Inhibitor of DNA Binding 4 HLH Protein (ID4), and Fibroblast Growth Factor Receptor 3 (FGFR3),[43] however, the quiescent SSC marker Undifferentiated Embryonic Cell Transcription Factor 1 (UTF1)[43] was upregulated 5±1-fold (**Figure 4**). The pan-germ cell marker DEAD-Box Helicase 4 (DDX4)[49] and was upregulated 7±1-fold (**Figure 4**), and Deleted in Azoospermia Like (DAZL), a broad translational regulator of genes associated with SSC proliferation, differentiation and survival,[43, 50] was likewise increased 12±1-fold (**Figure 4**).

**Figure 4:**
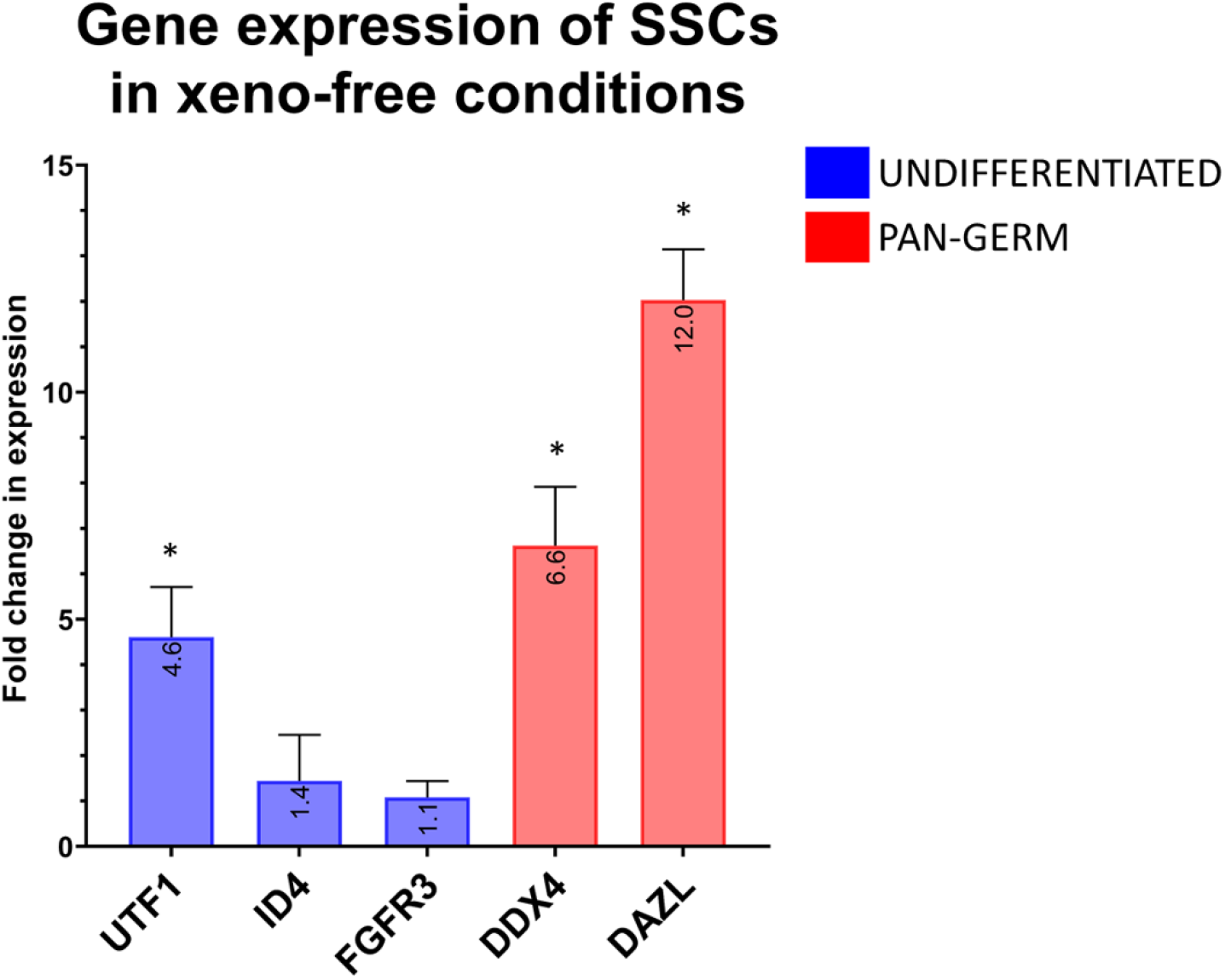
Gene expression of the long-term SSC cultures in xeno-free media compared to standard media. UTF1, ID4 and FGFR3 are quiescent SSC markers. DDX4 and DAZL are expressed by all germ cells. ^*^indicates a statistically significant difference compared to the standard media. **Abbreviations:** SSC = spermatogonial stem cell, UTF1 = Undifferentiated Embryonic Cell Transcription Factor 1, ID4 = Inhibitor Of DNA Binding 4 HLH Protein, FGFR3 = Fibroblast Growth Factor Receptor 3, DDX4 = DEAD-Box Helicase 4, DAZL = Deleted in Azoospermia-Like.

## 4.0 DISCUSSION

We show that the animal components FBS and BSA in standard SSC expansion media can be replaced by the growth factors PGD-2 and IGF-1 without loss of viability or expansion capability for at least 3 passages and 3 freeze-thaw cycles. Moreover, the cellular morphology of these cells begins to resemble the characteristic grape-like clusters seen when SSCs are cultured on mouse embryonic feeder (MEF) cells,[33] or in mixed testicular cultures,[48] suggesting a more natural phenotype. SSC populations in xeno-free or standard media were otherwise confirmed to share identical protein expression by immunocytochemistry analysis for well-known SSC markers. RNA expression was noted to increase in quiescent SSC and pan-germ markers, indicating that the xeno-free media was more supportive of SSC renewal than the standard media.

The most significant upregulation in gene expression was noted to be DAZL, a germ cell translational regulator that enhances gene expression by RNA binding and regulation of transcription factors and epigenetic regulators.[50, 51] It has a wide range of targets involved in processes from SSC proliferation and survival to differentiation. Observations in DAZL knockout mice suggest that one of its main functions is to enhance steady-state levels of SSC proliferation transcriptional regulatory factors,[51] suggesting that the xeno-free media promoted improved regulation of SSC proliferation.

There is currently one other reported xeno-free formulation for the expansion of SSCs,[52] however it relies on undefined human platelet lysate (hPL), which is a growth factor-rich alternative to FBS, collected from human platelets. While hPL is xeno-free, lot-to-lot variations are still a concern, as is the risk of pathogen contamination.[53, 54] Another limitation to the use of hPL is its ill-defined nature, since it introduces unknown factors to the cells, which can be detrimental to experimental design and analyses.

The effects of IGF-1 and PGD-2 on human germ cells are not well studied, however studies in mice suggest that IGF-1 acts in concert with GDNF and FGF-2 to promote complete cell cycle progression,[35] while PGD-2 supports gonocyte differentiation into SSCs and activates a quiescent state by repressing genes associated with the regulation of pluripotency, cell cycle arrest, and entry into meiosis.[36–38] Therefore, a possible explanation for their combined efficiency in promoting human SSC expansion is that together they regulate cell cycle progression while inhibiting pluripotency and pre-meiotic gene expression.

## 5.0 CONCLUSIONS AND FUTURE DIRECTIONS

This study shows that the growth factors PGD-2 and IGF-1 can replace the animal-derived components in established SSC expansion cell culture. This xeno-free, defined formulation can be used to standardize SSC *in vitro* culture, and removes the risk of pathogens and other unknown components.

Further investigation into the precise molecular mechanisms underlying the effectiveness of PDG-2 and IGF-1 will help to better define and inform future culture practices. Additionally, experiments using xenotransplant models could assess the capability of xeno-free SSCs to colonize the testicular niche and differentiate.

## 6.0 ACKNOWLEDGEMENTS

The authors would like to acknowledge the Vancouver Prostate Centre for their funding support and our collaborators at the Willerth Lab at the University of Victoria and the Lange Lab at the Vancouver Prostate Centre for their generous assistance.

## 8.0 DATA AVAILABILITY STATEMENT

The data that support the findings of this study are available from the corresponding author upon reasonable request.

## Footnotes

1 ACTA2: actin alpha 2, smooth muscle, BSA:bovine serum albumin, CREB:Clinical Research Ethics Boards, DAZL:deleted in azoospermia like, DDX4 -= DEAD-box helicase 4, EGF:epidermal growth factor, FBS:fetal bovine serum, FGF2:fibroblast growth factor 2, FGFR3:fibroblast growth factor receptor 3, GDNF:glial cell derived neurotrophic factor, GFRA1:GDNF family receptor alpha 1, GPR125:G protein-coupled receptor 125, hiPSC:human induced pluripotent stem cell, hPL:human platelet lysate, ID4:inhibitor of DNA binding 4, HLH protein, IGF-1:insulin-like growth factor 1, LIF:leukemia inhibitory factor, MEF:mouse embryonic feeder, OCT4:octamer binding protein 4, PGD-2:prostaglandin D2, PRM1:protamine 1, ROCK:Rho-Associated, Coiled-Coil Containing Protein Kinase, SOX9:SRY-box transcription factor 9, SSC:spermatogonial stem cell, SSEA4:stage specific embryonic antigen 4, STRA8:stimulated by retinoic acid 8, SYCP3:synaptonemal complex protein 3, THY1/CD90:Thy-1 Cell Surface Antigen, UTF1:undifferentiated embryonic cell transcription factor 1

